# Molecular dynamics simulation reveals that switchable combinations of β-sheets underlie the prion-like properties of α-synuclein amyloids

**DOI:** 10.1101/326462

**Authors:** Hiroki Otaki, Yuzuru Taguchi, Noriyuki Nishida

## Abstract

Diversity of prion strains is one of the most mysterious traits of prions because they are mere aggregates of abnormally-folded forms of single protein species, prion protein (PrP^Sc^), without genome. Although the strain-specific properties are hypothesized to be enciphered in the strain-specific structures of PrP^Sc^ instead of nucleotide genome, specifically what structure can code the information remains an enigma due to the incompatibility of PrP^Sc^ with structural analyses. Although the strain diversity was regarded as unique to prions, recently other disease-associated amyloids of α-synuclein (αSyn) or tau are also reported to have “strains”. As detailed structures of αSyn amyloid are already identified and the properties of mutant αSyn associated with familial Parkinson’s diseases, e.g. A53T, H50Q, and G51D, have been characterized, structure-phenotype relations of this type of amyloid could be investigated by using the αSyn amyloid as a model. Here we intensively investigated the mutant αSyn amyloids by molecular dynamics simulation to characterize influences of mutations on the structures of homo- or hetero-oligomer stacks of the amyloid. The simulations revealed directionality of the amyloid stack, remote effects of the mutations on distant β-sheets, existence of at least two switchable interfaces/amyloid cores, and distinct effects of hetero-oligomerization depending on mutation types. Collectively, those findings implied a possible mechanism of the strain diversity of the amyloids which have multiple in-register parallel β-sheets side-by-side, and support the view that their prion-like properties are inherent in the characteristic structures. We expect that the notion is also applicable to PrP^Sc^.

## Introduction

Prion disease is a unique group of neurodegenerative disorders which are caused by conformational abnormality of prion protein (PrP) and used to be mistaken as viral infections because of very high infectivity, diverse clinicopathological pictures suggestive of existence of many virus strains, strain-specific host ranges, and species barriers along with new-host adaptations [1]. Since a prion in reality consists solely of PrP in its abnormal isoform (PrP^Sc^) [1], the mechanism how mere aggregates of a single protein acquire the virus-like properties has been a long-standing challenge to the current biology. For example, regarding strain diversity of prions, an enormous body of evidences have indicated that strain-specific pathogenic information is enciphered in strain-specific structures of PrP^Sc^[2], but specifically what structures could code the information is unsolved yet, partly because detailed structures of PrP^Sc^ have not been identified due to its incompatibility with high-resolution structural analyses. Based on currently available data, PrP^Sc^ is hypothesized to be either in-register parallel β-sheet amyloid [3] or β-solenoid [4].

Many neurodegenerative diseases including Alzheimer’s disease (AD) and Parkinson’s disease (PD) are also caused by amyloid formation of their respective responsible proteins, e.g. Aβ peptide in AD, α-synuclein (αSyn) in PD, and tau protein in tauopathies. Interestingly, those disease-associated amyloids are often in-register parallel β-sheet amyloids [5][6][7][8] and have “prion-like” properties in common albeit lesser extents than those of authentic prions, suggesting a possibility that the properties are inherent in the in-register parallel β-sheet structures: for example, seeding efficiencies by an amyloid can be inefficient in heterologous reactions where substrates and template molecules are distinct in primary structures [9], and clinicopathological pictures of familial PD vary depending on mutations [10], reminiscent of species barriers and strain diversity of prions, respectively. Those similarities among the amyloids imply a possibility that knowledge from investigation on one amyloid is applicable to all the others.

If the prion-like properties are inherent in the structures of the in-register parallel β-sheet amyloids, structural investigation of those amyloids can provide a clue to elucidate the prion-like properties. Although they also share the similar difficulties in structural analyses as PrP^Sc^, high-resolution structures are available for some amyloids including Aβ peptide [6], brain-derived tau protein [8], and *in vitro*-formed αSyn fibrils [7], which were determined by solid-state NMR (ssNMR) and/or cryo-electronmicroscopy. We thought that molecular dynamics (MD) simulation of in-register parallel β-sheet amyloids could provide an insight into their intrinsic properties which are responsible for the prion-like properties. We were particularly interested in the αSyn amyloid (PDB ID 2n0a)[7], because of its relatively large size and the “Greek-key” conformation with multiple β-sheets intricately interacting one another. αSyn is a cytoplasmic protein consisting of 140 residues, well-known as the main component of Lewy bodies which are characteristically observed in Parkinson’s disease and dementia with Lewy bodies [11], or as the precursor of the “non-amyloid-β component” (NAC) fragments of amyloid plaques associated with AD [12]. While the physiological micelle-bound conformation has two large α-helixes in the N-terminal-side domain and disordered C-terminal-side domain [13], in α-synucleinopathies including PD, αSyn are found in amyloid forms: the conformations seem to be associated with the clinicopathological pictures, like prions [10]. The relations between the structures of the αSyn amyloids and the clinicopathological pictures are well demonstrated by the variation in clinical features among familial PD associated with mutations of αSyn, e.g. A53T, G51D and H50Q [10], although the molecular mechanisms how those mutations precipitate amyloid formation leading to development of the characteristic diseases are unsolved. αSyn is also suitable for *in-vitro* experiments because the recombinant proteins can be easily purified and form amyloid *in vitro* which exerts toxicity on cultured cells or animals [14][15]; those features enable efficient assessment of consistency between the *in silico* findings and the *in vitro* behaviors of the protein. We therefore utilized αSyn for investigation of general properties of in-register parallel β-sheet amyloids, and as a potential surrogate model of PrP^Sc^. Since the detailed structures of the αSyn amyloid was available, we tested the validity of the hypothesis that the prion-like properties are inherent in the in-register parallel β-sheet structures in *in silico* settings.

The Greek-key conformation of αSyn amyloid has been analyzed in many *in silico* analyses including MD simulation. For example, Xu *et al.* assessed influences of pathogenic mutations on the amyloid [16], and postulated that the mutations switch preferred amyloid conformations of the substrate αSyn between the Greek-key conformation and a superpleated β-structure, whose distribution patterns of β-sheets were mutually rather distinct. Here, we devised a few points in MD simulation of αSyn amyloids for reliable and informative results. First, we used a ten-molecule stack of αSyn for MD simulation to characterize behaviors of molecules at different positions/layers in the stack: the molecules at the both ends of the stack, i.e. the Chains A and J (**Fig 1A, right panel**), are predictable to be most unstable, while in contrast those in the middle of the stack would be stable; then how the stability of the peptides transit from the middle to the end? We thought that ten layers would be thick enough to address the question. Second, we set the sampling time length at 400 ns per run for reliable evaluation of the stabilities of amyloid stacks and the influences of mutations on the structures; besides, each simulation was repeated three to five times for statistic evaluation to reduce influences of stochastic factors. Moreover, since most of the mutant αSyn-associated familial PD patients are heterozygous for the mutant allele, we analyzed hetero-oligomer A53T amyloids, which combine A53T mutant and wild-type αSyn at various ratios, to characterize the influences of the mutant molecules on the wild-type in the same stack.

**Fig 1.**
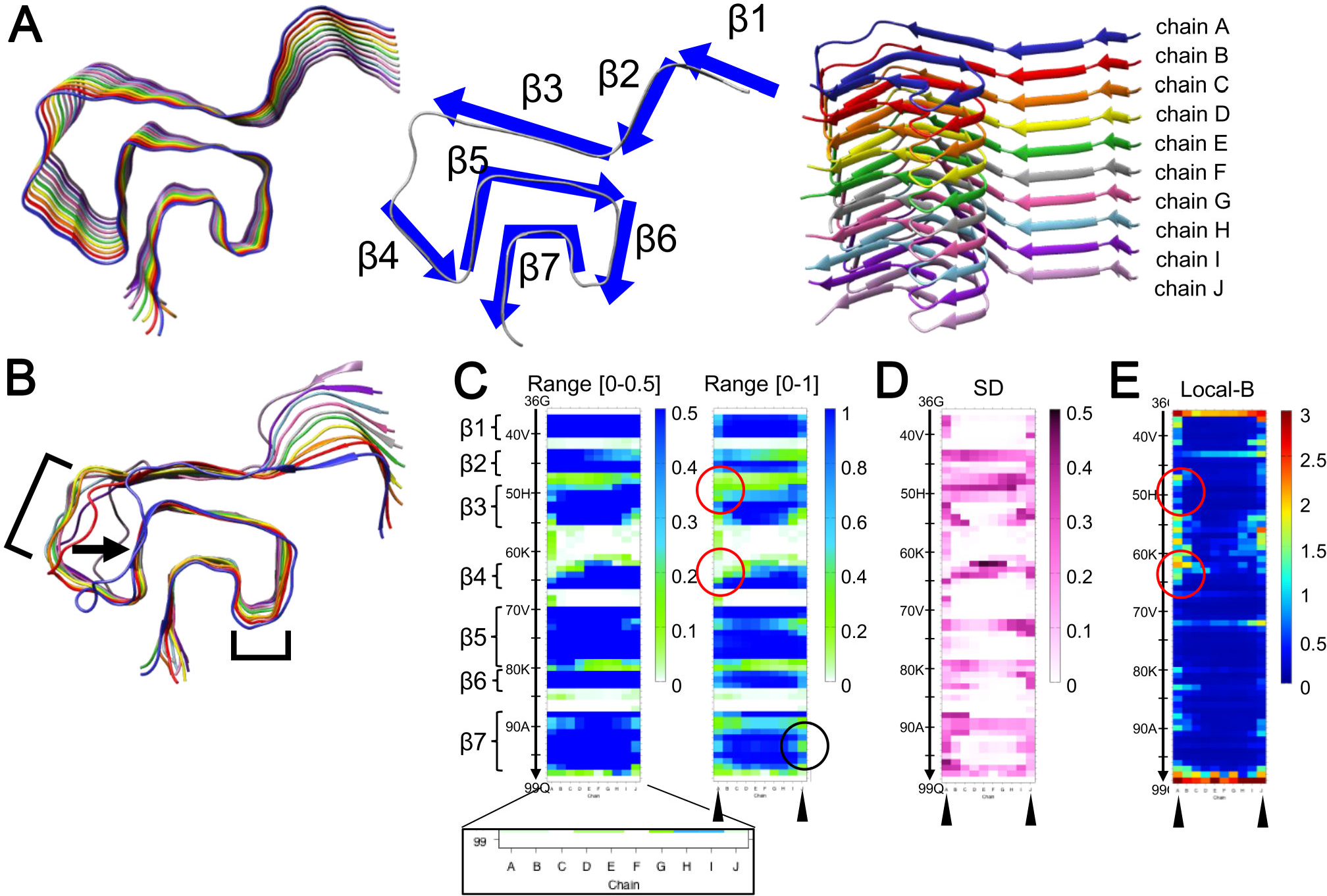
The Molecular dynamics (MD) simulation of the wild-type αSyn(WT) amyloid. **A.** The starting “Greek-key” conformations of the αSyn amyloid [PDB ID: 2n0a (Tuttle *et al.,* 2016)] for the MD simulation: the left panel show the top view of the amyloid stack. For simplicity, the β-sheets are numbered from β1 to β7 (middle panel) and the ten chains are referred to as the Chain A to J, from top to bottom (right panel). **B.** A representative image of WT amyloids after MD simulation for 400 ns, with the Chain A front side. Note that the stack-end molecules, Chains A and J, are substantially disordered particularly in the β3-β4 regions (arrow), while the Chains B-l are rather stable. All the images of the five independent runs are presented in the Supplementary information **(Suppl Fig S2)**. **C-E.** Heat maps illustrating average β-sheet propensities **(C)**, standard deviations (SD) of the β-sheet propensities **(D)**, and the backbone local B-factor (Local-B) values **(E)**, respectively. The values are based on five independent runs. The vertical axis represent residues from Gly36 to Gln99, and the horizontal axis corresponds to the Chain A to J (inset). **C.** Heatmaps of β-sheet propensity with value ranges 0-0.5 (left) and 0-1 (right). Note that the regions with high β-sheet propensities correspond to β1 to β7, and the stack-end molecules have relatively low values (arrowheads). Interestingly, color distributions were distinct between the Chain A and the Chain J sides (red circles). **D.** The heatmap of SD of the β-sheet propensity revealed that the color distributions were distinct between the Chain A and J sides, particularly in the β5 and β7, with a tendency to be high at residues with relatively low β-sheet propensities. **E.** The heatmap of the backbone local B-factors show relatively high values in the loop region between β3 and β4, and that the Chains A and J have distinct color distributions (arrowheads).

To our surprise, the relatively small and simple αSyn amyloids exhibited behaviors reminiscent of characteristic properties of prions, particularly strain diversity and species barriers. The results seemed to support our view that the prion-like properties of amyloids are inherent in the in-register parallel β-sheet structure, a rather singular status where many molecules take the same conformations and intimately interact. We expect that the knowledge is applicable to other disease-associated amyloids as well, to help address their unsolved problems derived from in-register parallel β-sheet structures.

## Material & Methods

### Modeling Structure

We used the Greek-key conformation of the αSyn amyloid (PDB ID 2n0a) for the starter. Since our interest was in the general properties of in-register parallel β-sheet amyloids rather than αSyn itself, we chose to truncate the N- and the C-terminal disordered regions, namely 1-35 and 100-140, to reduce calculation burdens. Then, to minimize the electrostatic interactions at the N- and C-termini, we modified them with acetylation and methylation using AmberTools16 [17], respectively. We ascertained that behaviors of the “capped 36-99” were not very different from those of either “uncapped 36-99” or “capped 20-110” (data not shown). For simplicity of description, we numbered the β-sheets from β1 to β7 (**Fig 1A, middle panel**), and refer to each layer from top to bottom as the Chain A to J (**Fig 1A, right panel**). For modeling of the mutant αSyn including A53T, G51D, and H50Q, we used SCWRL4 [18]. Modeled amyloid was solvated with a rhombic dodecahedron water box with minimum wall distance of 12 Å. Na^+^ and Cl^−^ ions were randomly placed to neutralize the system and yield a net concentration of 150 mM NaCl. Protonation status of the histidine at the codon 50 was fixated as N^δ^-H tautomer (HID form in AMBER naming convention) in all the simulations.

### MD simulation

For all the MD simulation, we used GROMACS software [19] with AMBER ff99SB-ILDN force fields [20] for the αSyn amyloids and the TIP3P water model [21]. The system was minimized for 5,000 steps with steepest descent method, followed by 2,000 steps with conjugate gradient method. During the minimization, heavy atoms of the amyloids were restrained with a harmonic potential with the force constant of 10.0 kcal/mol•Å^2^. After the minimization, the temperature of the system was increased from 0 to 310 K during 1 ns simulation with the restraints. Next, 1 ns of equilibration run was performed by gradually reducing the restraints from 10.0 kcal/mol•Å^2^ to zero and subsequent equilibration was performed in the NPT ensemble for 2 ns at 310 K and 1 bar. Production runs were carried out for 400 ns in the NPT ensemble at 310 K and 1 bar (**Suppl Fig S1A**). We used velocity-rescaling scheme [22] and Berendsen scheme [23] for thermostat and barostat, respectively. The LINCS algorithm [24] was used to constrain all bonds with hydrogen atoms, allowing the use of 2 fs time step. Electrostatic interactions were calculated with the Particle-mesh Ewald method [25]. The cutoff length was set to 12 Å for Coulomb and van der Waals interactions. The Verlet cutoff scheme [26] was used for neighbor searching. We have carried out 5 MD simulations for the wild-type amyloid and the homo-oligomer A53T amyloid and 3 MD simulations for the other mutants. For each production run, trajectory snapshots were saved at every 10 ps.

### Analyses

For the analysis of the MD simulations, the first 100ns of each trajectory was discarded except for calculation of root-mean-square deviation (RMSD). Thus, 30,001 snapshots (trajectory from 100 to 400ns) were used for the analysis. The RMSD for backbone atoms of every chain was calculated using *gmx rms* program in GROMACS package. The ssNMR structure was used as the reference structure. Although we first tried ordinary root-mean-square fluctuation (RMSF) using *gmx rmsf* program in GROMACS for evaluation of the structural stabilities of each peptides, it proved not very suitable for the αSyn amyloid because the deviations in the β1-β2 regions were so large as to mask the deviations in the other regions (**Supplementary Movie**). We therefore decided to utilize the local backbone B-factor suggested by Fuchs et al [27]. The local backbone B-factor is calculated via multiple local alignments to all backbone atoms including hydrogen of the residue of interest instead of global alignment for usual RMSF. The local backbone B-factor is evaluated as the mass-weighted B-factor of the atom groups used for the alignment. We also use the ssNMR structure as the reference for alignment and *cpptraj* [28] in AmberTools for calculation. Hydrogen bond pattern was characterized according to their occupancy in the trajectory with geometric criteria using *H-bond* plugin in VMD [29]. A hydrogen bond is assigned if the distance between donor (D) and acceptor (A) is less than 3.2 Å and the angle A-D-H is less than 30°. The secondary structure content during the simulations was calculated using DSSP program [30][31] and *gmx do_dssp* program in GROMACS. Hydrophobic contact was analyzed using *PyInteraph* [32]. The hydrophobic contact is assigned if the distance between the centers of mass of side-chains is less than 5 Å [33]. The results of the hydrogen bond analysis and hydrophobic contact analysis were visualized using Cytoscape [34]. All molecular structures for αSyn amyloids were drawn using UCSF Chimera [35] and simulation movie was generated using VMD [29].

### Short MD runs

We noticed that certain effects of mutations were already present at the very starting point of the product run of MD simulation and would remain unchanged or even accentuated over the subsequent sampling time, suggesting that the effects manifest in the pre-run phase. In such cases, we exploited it and repeated short MD simulation 50 times to exclude a possibility that the apparent effects of the mutation actually just resulted from probability fluctuation. We carried out 5 ns of production runs from the common minimized structure according to the same procedure as the 400 ns simulations (**Suppl Fig S5A**).

### Statistical analyses

For analysis of statistical differences between two groups, we used the two-tailed Student t-test of the Excel2013 (Microsoft Corporation, Redmond, WA). A p-value less than 0.05 was considered as statistically significant.

## Results

### MD simulation of wild-type αSyn amyloid

First, we ascertained stability of the stack of wild-type αSyn [αSyn(WT)] amyloid in the MD-simulation environment (**Fig 1B**). The αSyn(WT) amyloid molecule remained mostly stable over 400 ns of sampling time albeit relative instability in the two loop regions of the stack-end molecules, the Chains A and J, as anticipated (**Fig 1B, brackets**). Interestingly, the Chains A and J showed different degrees of instability: the loop encompassing the residues 56-62 (**Fig 1B, larger bracket**) and the adjacent β-strands, β3 and β4, of the Chain A were more disordered than those on the Chain J side (**Fig 1B, arrow**). Stabilities of the stack-end molecules seemed varied also between the five runs (**Suppl Fig S2A**).

Analysis of β-sheet propensity revealed more details of the differences between the chains. The stack-end Chain A and J had the most unstable β-sheets as indicated by the low average β-propensity values (**Fig 1C**) and large standard deviation (SD) values (**Fig 1D**). In addition to the β-sheet propensity analysis, we also analyzed local B-factors to quantify regional mobility [36] and found them consistent with the images of MD simulation (**Fig 1E**). The Chains C-H showed constantly high β-propensity and low SD values in most β-sheets, corroborating structural stability of the middle layers. The heatmaps also corroborated differences in structural stabilities between Chains A and J: The average β-sheet propensities of the β3 and β4 regions were lower in the Chain A than in the Chain J (**Fig 1C, red circles**) along with higher B-factor values (**Fig 1E, red circles**), whereas the β7 region of the Chain J showed lower β-propensity values than the Chain A (**Fig 1C, black circle**). These distribution patterns of high β-sheet propensity could be important determinants of properties of the amyloid because the stack-end molecules are theoretically the immediate interfaces for the amyloid to interact with other cognate amyloidogenic proteins or non-amyloidogenic proteins. We therefore mainly focus on the stack-end molecules, the Chains A, B, I and J, from here on.

### MD simulation of homo- and hetero-oligomer A53T amyloid

After the characterization of behaviors of the αSyn(WT) amyloid (WT amyloid) in the MD simulation, we assessed influences of a representative familial PD-associated substitution mutation A53T [10]. The homo-oligomer (homo-A53T) amyloid of the mutant αSyn with A53T substitution [αSyn(A53T)] apparently behaved similarly to WT amyloid, except for a tendency to lose β-sheet structures in β7 of the Chains A and B (**Fig 2A, arrow**). Analysis of β-sheet propensity and local B-factor values confirmed the remote effects of the A53T substitution on the β7 region, which is too distant for the residue 53 to directly interact: average β-sheet propensity values of homo-A53T amyloid were obviously lower in the β7 region of the Chains A, B, I, and J than those of WT (**Fig 2B, red circle and bracket**), along with higher SD values suggesting poorer reproduction of β-sheet structure (**Fig 2C, bracket**) and larger local B-factor particularly in the Chain A and B reflecting local structural instability (**Fig 2D, red circle**). The A53T substitution reduced β-sheet propensity values of the β7 regions of the Chain A and B similarly (**Fig 2E**), and the differences from those of WT in five independent runs were statistically significant (**Fig 2F, right column**). The propensity values of β7 of the Chain J also tended to be lower than those of WT, but the differences were not significant. Although less prominent than β7, the A53T substitution also affected the β5 region: the β-sheet propensities of the residue 74 of the Chain A, B, I and J tended to be lower in the homo-A53T than in WT, while those of the residue 72 oppositely tended to be higher (**Fig 2F, left column**). Those alterations in the β5 prompted us to analyze hydrogen bonds and hydrophobic effects of the amyloid, because β3 and β5 were close enough to directly interact (**Fig 3**). As expected, hydrogen bonds were newly formed between the substituted Thr53 and Gly73 or Val74 (**Fig 3A, red lines**), which reminisced the speculative model by Rodriguez *et al.* [37]. A noteworthy fact is that they were “inter-layer” bonds bridging a Thr53 with the Gly73 of one-upper layer, e.g. Thr53 of the Chain B and Gly73 of the Chain A (**Suppl Fig S3**). To the contrary, hydrophobic effects between Ala53 and Val74 in the WT are absent in the homo-A53T amyloid (**Fig 3B**). The diagrams also revealed interactions between β5 and β7, e.g. hydrogen bonds between Gln79 and Ala89 or hydrophobic effects connecting Val77 and Ala90, and those interactions could affect the structures of β7 when β5 is affected by the A53T mutation.

**Fig 2.**
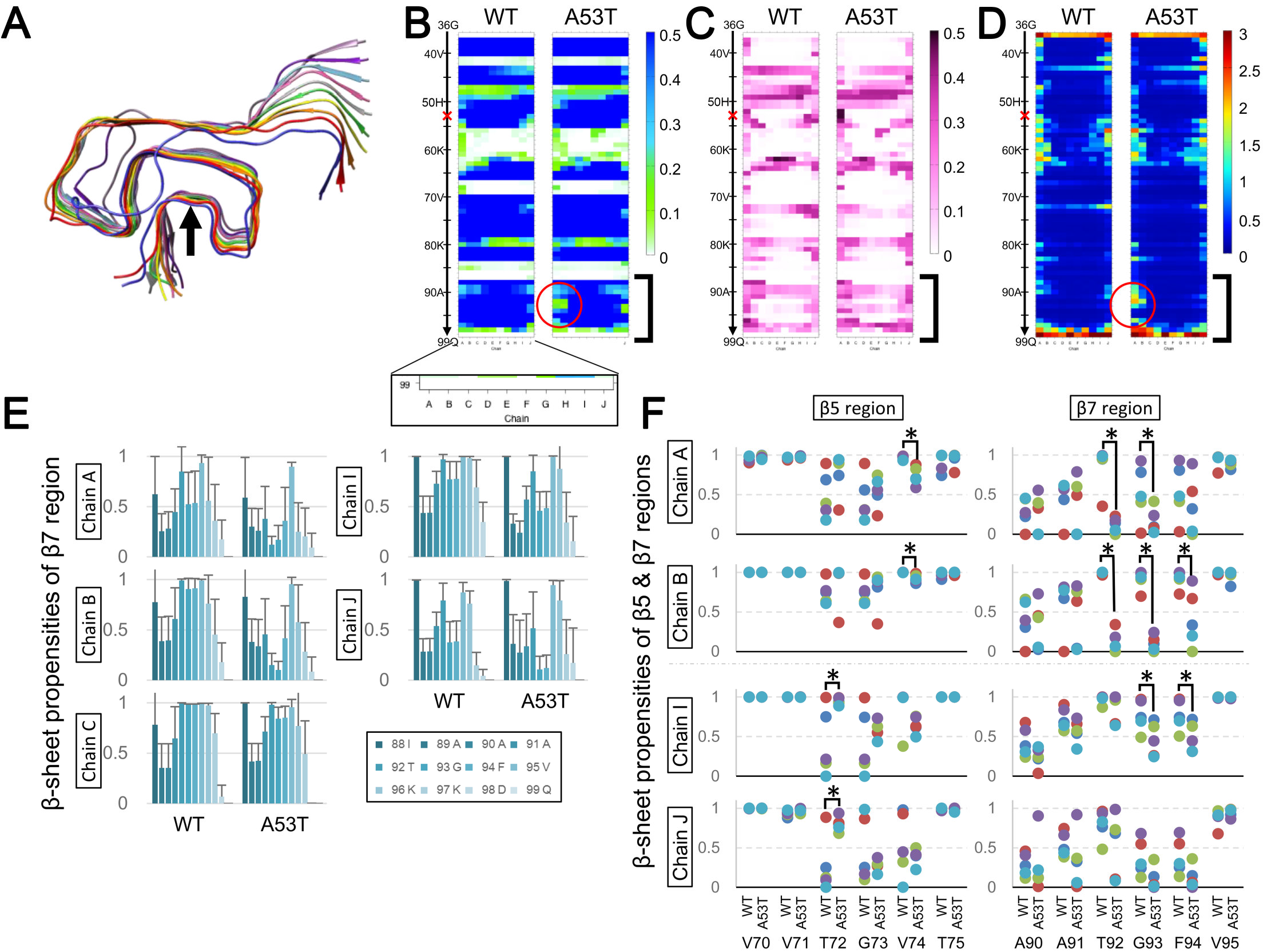
The homo-A53T amyloid is uniquely unstable in the β7 region. **A.** A representative image of the homo-oligomer (homo-A53T) amyloid of αSyn(A53T) after 400 ns of the sampling time. All the images of the five independent runs are presented in the **Supplementary Figure (Suppl Fig S2).** The β7 region of the Chain A and B are not in the β-sheet form (arrow). **B-D.** Comparison of heatmaps of average β-sheet propensities (**B**), SD of the β-propensities (**C**), and backbone local B-factors (**D**) between WT and homo-A53T amyloids (the left and the right of each pair, respectively). The values are based on five independent runs. The vertical axis represent residues from Gly36 to Gln99, and the horizontal axis corresponds to the Chain A to J (inset). The red cross on the vertical axis indicates the position of the A53T substitution mutation. **B.** Comparison of β-sheet propensities of WT and homo-A53T amyloids revealed that the β-sheet propensities of homo-A53T are substantially lower than the WT amyloid in the β7 region of the Chains A and B (red circle, bracket). **C.** SD of the homo-A53T amyloids are also higher in the β7 region than that of WT (bracket), suggesting poorer reproducibility of the regional structures. **D.** The local B-factors are higher in β7 of the Chain A and B of homo-A53T amyloid (red circle), i.e. higher mobility of the chains in the region, consistent as local structural instability. **E**. Bar graphs of β-sheet propensities of the β7 regions (residues 88-99) of the stack-end Chains A, B, C, I and J, comparing the WT and the homo-A53T amyloids. The bars and the error bars represent the average values and the SD of five independent runs, respectively. **F.** Dot plots of β-propensities of the β5 regions (residues 70-75) and the β7 region (residues 90-95) of the stack-end Chains A, B, I and J, comparing the WT and the homo-A53T amyloids. Asterisks, p < 0.05. Note that the β-propensity values in the β7 of the Chain A and B of homo-A53T are similarly lower than those of WT.

**Fig 3.**
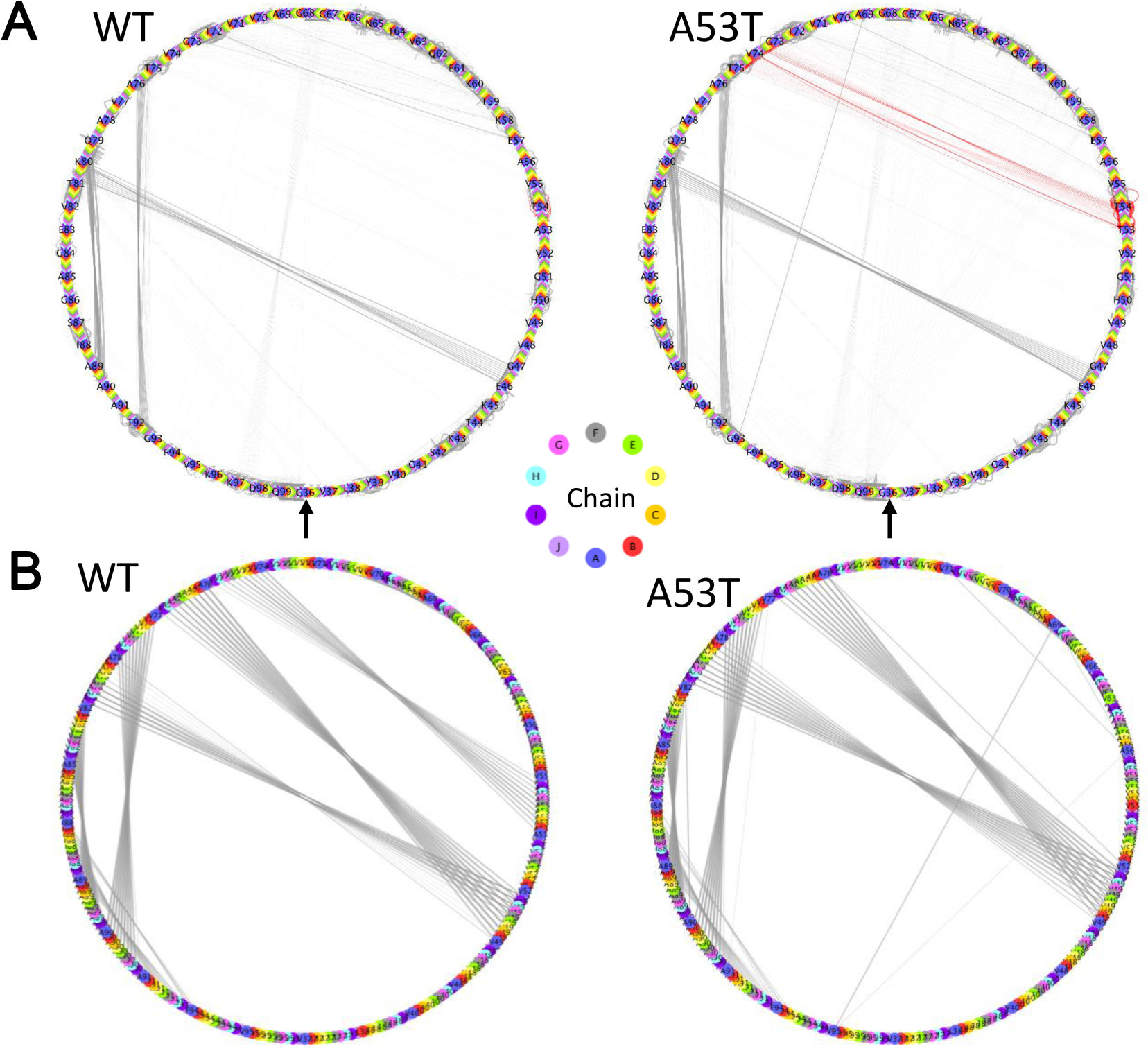
Hydrogen Bond and Hydrophobic Networks: comparison between WT and homo-A53T amyloids. **A.** Diagrams illustrating hydrogen bond networks of the WT and homo-A53T amyloids. Color code for each chain is presented between them. Residue 35-99 of αSyn are arranged from the bottom (arrow) of the circles in a counterclockwise fashion, i.e. 36(A,B…J), 37(A-J), … 99(A-J). Note that there are unique hydrogen bonds in the homo-A53T amyloid, e.g. T53-V74 and A69-G93, and the bonds involving A53T are high-lighted in red. A magnified view of a part of a circle is presented **(Suppl Fig S3B)** to show how the chains are arranged in the diagram. **B.** Diagrams illustrating hydrophobic effects in the αSyn amyloids. Unlike the **Fig 3A**, the circles contain only residues contributing to hydrophobic effects. Color codes for each chain is the same as above. Note that a bundle of hydrophobic effects between the residues 53 and 74 in the WT amyloid is absent in the A53T amyloid.

Given the discernible differences between WT- and homo-A53T amyloids, we analyzed hetero-oligomer amyloids containing both αSyn(WT) and αSyn(A53T) at various ratios. Since most familial-PD patients are heterozygous for the pathogenic mutations, expressing both wild-type and mutant αSyn, this could appropriately recapitulate the *in vivo* circumstances. Which phenotype predominates when they are mixed in a stack? To address the question, we analyzed the following hetero-oligomer A53T (hetero-A53T) amyloids: αSyn(A53T) at (1) Chains A and J (AJ), (2) Chains A, B and J (ABJ), (3) Chain A only, (4) Chain J only, (5) Chains A and B (AB), (6) Chains I and J (IJ), (7) Chains A to C (ABC), (8) Chains H to J (HIJ), and (9) Chains A, B, C, D and E (ABCDE). As a result, all the hetero-A53T amyloids seemed more similar to WT amyloid than homo-A53T in terms of the β-sheet propensity values of β7 in the Chains A and B (**Fig 4A and 4B**), i.e. the unique instability of homo-A53T amyloid was suppressed by the hetero-oligomerization. The backbone Local B-factors were also consistent with the results of the β-sheet propensity values (**Suppl Fig S4**). Interestingly, hetero-A53T amyloids with αSyn(A53T) on the Chain-A side, e.g. AB-, ABC- and ABCDE-amyloid, tended to have higher β-sheet propensity values in β3 of the Chain A and B than WT particularly in the residues 52-54 (**Fig 4C**), suggesting that they are not simple phenocopies of the WT amyloid. The newly-formed hydrogen bonds between Thr53 and Val52 may contribute to this stabilization of β3 (**Suppl Fig S3**). Besides, characteristics of the homo-A53T were also hinted in β-sheet propensity values at the residue 74 of β5, particularly in those with αSyn(A53T) clustered on the Chain-A side, again (**Fig 4D, compare with “A53T”**). To our surprise, even a single mutant in an amyloid stack could affect properties of all the chains in the stack, as seen in the small variations in the β-sheet propensity values of Gly93-Phe94 of the Chain A in the “J only” (**Fig 4E**). “IJ” and “HIJ” showed the similar tendencies. Possibly, he numerous hydrogen bonds and salt bridges between the chains facilitate conduction of motions over a long range across the stack, so that all the chains share the motion irrespective of the primary structures.

**Fig 4.**
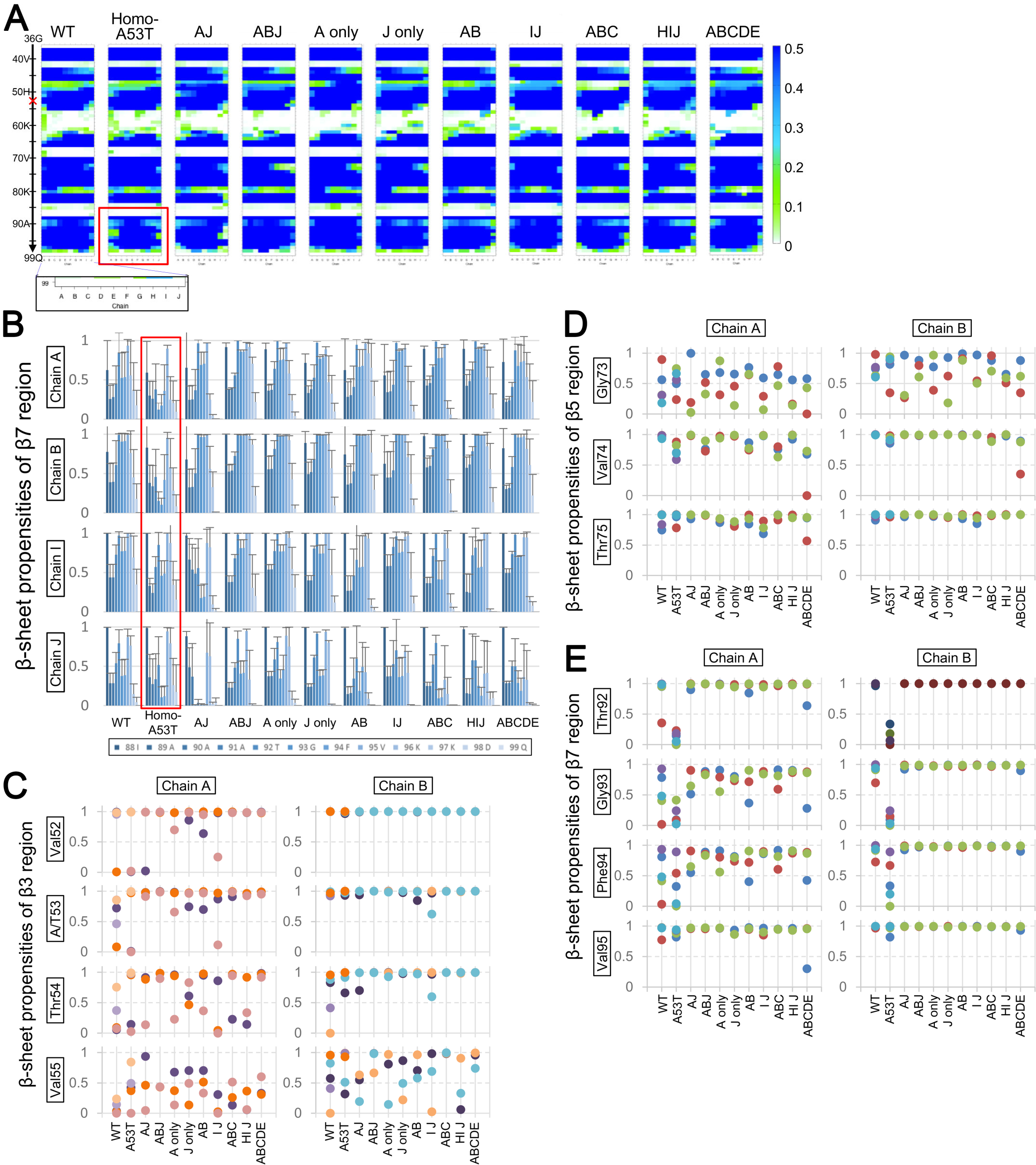
The hetero-oligomers of αSyn(WT) and αSyn(A53T) are distinct from either homo-A53T or WT amyloid. **A.** Heatmaps of average β-sheet propensities comparing WT, homo-A53T, and various hetero-A53T amyloids. The values of various hetero-A53T are averages of three independent runs, while those of WT and homo-A53T are averages of five runs. Note that the β-propensity profiles of the hetero-A53T amyloids are more similar to that of WT regarding the β7 region, while those of homo-A53T are uniquely low (red box). The red cross indicates the position of the A53T substitution mutation. **B.** Bar graphs of the β-sheet propensities of the β7 regions of the stack-end Chains A, B, I and J, comparing WT, homo-A53T (red box), and various hetero-A53T amyloids. The bars and the error bars represent the average values and the SD of five (for WT and homo-A53T) or three (for hetero-A53T) independent runs, respectively. **C-E.** Dot plots of the β-sheet propensities in β3 (residues 52-55) (**C**), β5 (73-75) (**D**) and β7 (92-95) **(E)** regions of the stack-end Chains A and B comparing WT, homo-A53T and various hetero-A53T amyloids. Run numbers are the same as above.

### Short MD runs confirm the singularity of the homo-A53T amyloid

The low β-sheet propensity of β7 in homo-A53T amyloid was likely to be a unique feature attributable to the A53T substitution. However, there still was a possibility that they just resulted from probability fluctuation. As we noticed that the β7 region of the homo-A53T was already disordered at the very starting point and would remain unchanged or even accentuated over the subsequent sampling time, we attempted to address the possibility by repeating short MD simulation runs (**Suppl Fig S5A**). As anticipated, the diminution of β-sheet propensities in the β7 region of homo-A53T amyloid reproducibly occurred in the very initial step of the simulations (**Fig 5A, “0 ns”**). As the hetero-A53T(ABCDE) was similar to WT amyloid as described, we increased the ratio of αSyn(A53T) and performed short-MD simulations to identify when the characteristics of the homo-A53T emerge (**Fig 5B**). Surprisingly, none of the hetero-A53T amyloids was similar to the homo-A53T amyloid, including even the one with the mutant only at the Chain J, i.e. hetero-A53T(A-I) (**Fig 5B; Suppl Fig S5B, S6**).

**Fig 5.**
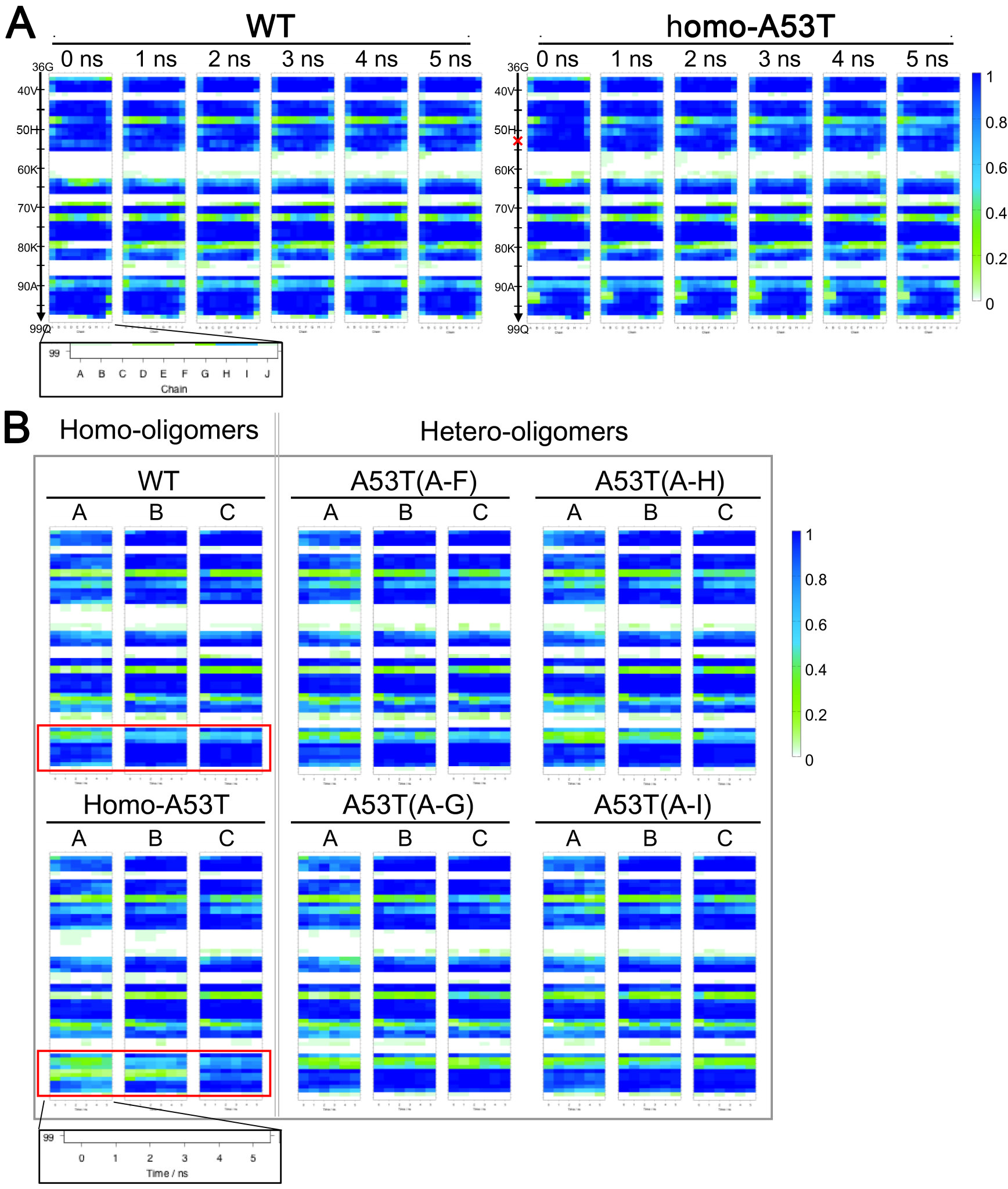
The unique loss of β-sheet in the β7 region of the homo-A53T amyloid occurs in the very early phase. **A.** Heatmaps of β-sheet propensities calculated from 50 runs of the short-MD simulations, comparing those of WT and homo-A53T amyloids. Note that the low β-propensity in the β7 of the homo-A53T amyloid is already present at the point of “0 ns”. The vertical axis represents the residues from 36 to 99, and the horizontal axis represents the Chain A-J (inset). The red cross indicates the position of the A53T substitution mutation. **B.** Heatmaps of β-sheet propensities calculated from the short-MD simulations, comparing those of homo-oligomer amyloids and the hetero-oligomer amyloids in which αSyn(A53T) is the majority. Each heatmap represent longitudinal fluctuations in the average β-sheet propensity values over the production run of 5 ns for the Chain A, B or C. The horizontal axis represents the time of production run (inset). Not only the Chains A and B of the homo-A53T, even the Chain C show lower β-sheet propensities in the β7 region than those of WT (compare the red boxes). The low β-sheet propensities in the β7 region at the very early phase of simulation are unique to the homo-A53T amyloid and the hetero-oligomers could not acquire the phenotype, even when the ratio of αSyn(A53T) in the stack was increased up to the point only the Chain J is αSyn(WT) [“A53T(A-I)”].

### MD simulation of homo-H50Q and homo-G51D amyloids

Unlike the homo-A53T amyloid, the homo-oligomer (homo-H50Q) amyloid of αSyn with H50Q substitution [αSyn(H50Q)] was rather similar to the WT amyloid in terms of apparent stability of β-sheets on the Chain-A side (**Fig 6A**). The β-sheet propensity and the local B-factor values were also similar to those of WT, although the β7 region of the Chain J seemed slightly unstable than that of WT (**Fig 6B and 6C, bracket**). On the other hand, the local B-factors in the β3 region were smaller than those of WT (**Fig 6C, red box**). Characteristically, there were inter-chain hydrogen-bond formation between the substituted Gln50, i.e. so-called a glutamine ladder (**Suppl Fig S7**), possibly contributing to the stability of β3 (**Fig 6E, left column**). Unlike A53T, H50Q did not affect β-sheet propensity values of β5, and those of the β7 region of the Chain J were only slightly smaller than those of WT (**Fig 6D and 6E, right columns**). As for the hetero-oligomer amyloids, while hetero-H50Q(ABC) amyloid was very similar to homo-H50Q amyloid or WT (**Suppl Fig S8**), hetero-H50Q(ABCDE) amyloid occasionally showed substantial deformity in β6 and β7 particularly on the Chain-J side (**Suppl Fig S9A, arrows**), which is also reflected in the low β-sheet propensity values of β7 (**Suppl Fig S9B, red box**). The present study indicates that αSyn(H50Q) can behave like αSyn(WT), or even advantageously in the amyloid formation process because of the glutamine ladder. Regardless, the age of onset of the H50Q-associated familial PD is relatively late [10] and, besides, pathogenicity of H50Q mutation itself is recently questioned [38]. Possibly, destabilization by hetero-oligomerization with αSyn(WT) might be responsible.

**Fig 6.**
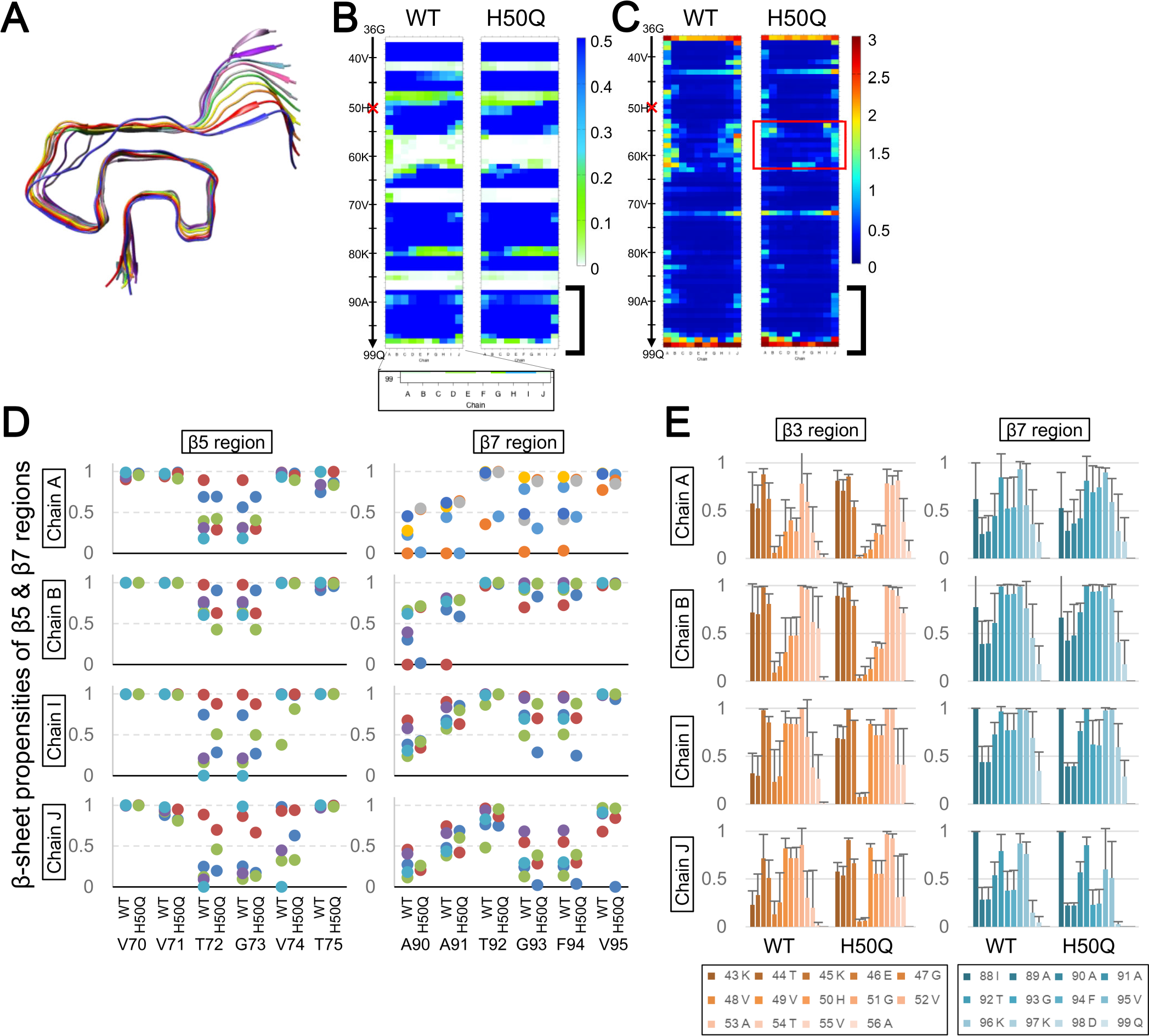
Homo-H50Q amyloid is very similar to the WT amyloid. **A.** A representative image of the homo-H50Q amyloid after 400 ns of sampling time. All the images from the three independent runs are presented in the **Supplementary Figure (Suppl Fig S7)**. **B-C.** Comparison of heatmaps of average β-sheet propensities (**B**) and backbone local B-factors (**C**) between WT and homo-H50Q amyloids. The values are based on five runs for WT, and three for homo-H50Q. The vertical axis represent residues from Gly36 to Gln99, and the horizontal axis corresponds to the Chain A to J (inset). The red cross on the vertical axis indicates the position of the H50Q substitution mutation. **D.** Dot plots of the β-sheet propensities in β5 (70-75) and β7 (90-95) regions of the stack-end Chains A, B, I and J, comparing WT and homo-H50Q amyloids. Run numbers are the same as above. **E.** Bar graphs of β-sheet propensities of β3 (residues 43-56) and β7 (88-99) of the stack-end Chains A, B, I and J, comparing the WT and the homo-H50Q amyloids. The bars and the error bars represent the average values and SD, respectively. The values are based on three runs for homo-H50Q and five runs for WT.

In MD simulation of αSyn amyloid with G51D substitution [αSyn(G51D)], the β3 region of homo-oligomer (homo-G51D) amyloid was severely disordered, with the peptides dissociating from one another (**Fig 7A, arrow; Fig 7B, 7D & 7E, red boxes**). The unique influence of the mutation was predictable because the substituted aspartate residues would align side-by-side with short intervals in the in-register parallel β-sheets and their negative charges would make the peptides mutually repel. In contrast to the severely disordered β3, the regions encompassing β5 to β7 seemed as stable as those of the WT amyloid (**Fig 7B, 7D & 7E, black boxes**). On the Chain-A side, the β-sheet propensities of the β7 regions were only slightly lower than those of the WT amyloid (**Fig 7E and 7F, right column**), while they tended to be higher than those of WT on the Chain-J side, although not statistically significant. The β5 region of the homo-G51D showed much lower β-sheet propensities at the residues 72 and 73 than those of WT (**Fig 7F, right column**).

**Fig 7.**
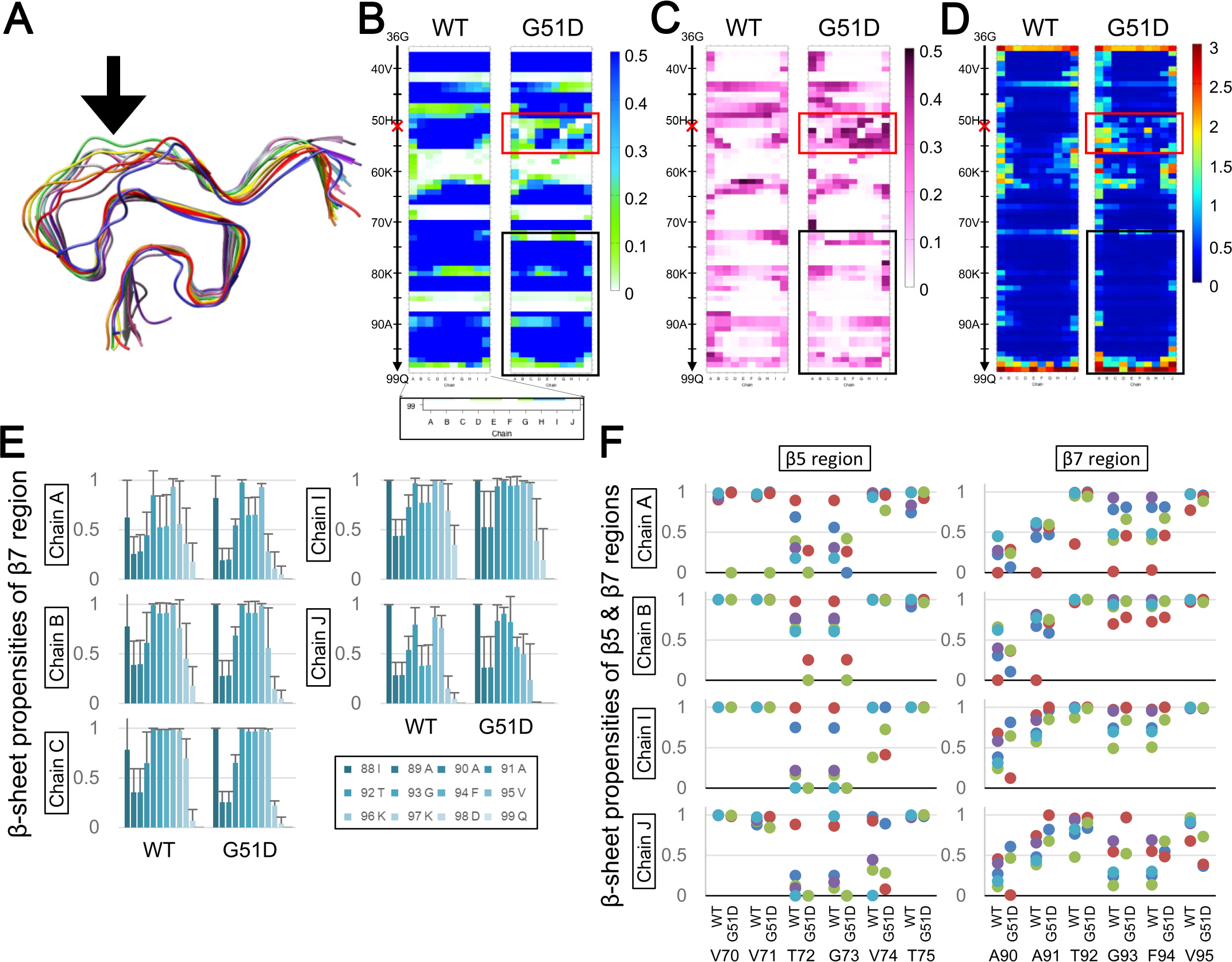
The homo-G51D amyloid has severely disordered β3 and stable β7. **A.** A representative image of the homo-G51D amyloid after 400 ns of sampling time. All the images from the three independent runs are presented in the **Supplementary Figure (Suppl Fig S11A)**. **B-D.** Comparison of heatmaps of average β-sheet propensities (**B**), SD of the β-sheet propensity values (**C**) and backbone local B-factors (**D**) between WT and homo-G51D amyloids. The values of the homo-G51D are based on three independent runs. The vertical axis represent residues from Gly36 to Gln99, and the horizontal axis corresponds to the Chain A to J (inset). The red cross on the vertical axis indicates the position of the G51D substitution mutation. The parameters corroborate that the β3 region of homo-G51D is severely deranged (red box). On the other hand, the β7 region shows comparable β-sheet propensity profile along with relatively low SD values in the region (black box). **E.** Bar graphs of β-sheet propensities of the β7 regions (residues 88-99) of the stack-end Chains A, B, C, I and J, comparing the WT and the homo-G51D amyloids. The bars and the error bars represent the average values and SD, respectively. The run numbers are the same as above. **F.** Dot plots of β-propensities of the β5 regions (residues 70-75) and the β7 region (residues 90-95) of the stack-end Chains A, B, I and J, comparing the WT and the homo-G51D amyloids.

### MD simulation of homo-(G68E+V95G), homo-S87N & homo-(A53T+S87N) amyloids

Species barriers are one of characteristic features of prions. Efficiencies of cross-seeding reactions, i.e. seeding reactions between amyloidogenic proteins with non-identical primary structures, can be equivalent to the species barriers in interspecies transmissions of prions. We were therefore interested in influences of mutations which are reported to greatly affect the cross-seeding efficiencies of αSyn amyloids, derived from ovine αSyn or mouse αSyn [39][9].

Human αSyn mutants with either of the two ovine residues, G68E or V95G, are not cross-seeded by fibrils of αSyn(WT) [39]: Gly68 and Val95 were located in the turn between β4 and β5 and at the C-terminal part of β7, respectively. Interestingly, not only the cross-seeding by human αSyn fibrils, fibril formation of sheep αSyn itself seems to be rather inefficient. In MD simulation of the homo-oligomer amyloid combining both G68E and V95G [homo-(G68E+V95E)], the global structures were unexpectedly stable (**Fig 8A, left panel**). Nevertheless, certain signs of incompatibility could be recognized, like reduced β-sheet propensity values in the distal β7 of the Chain A and B, and β4 of the Chain A (**Fig 8B, black circles**), accompanied by larger local B-factors suggesting local structural instabilities (**Fig 8C, black circles**).

**Fig 8.**
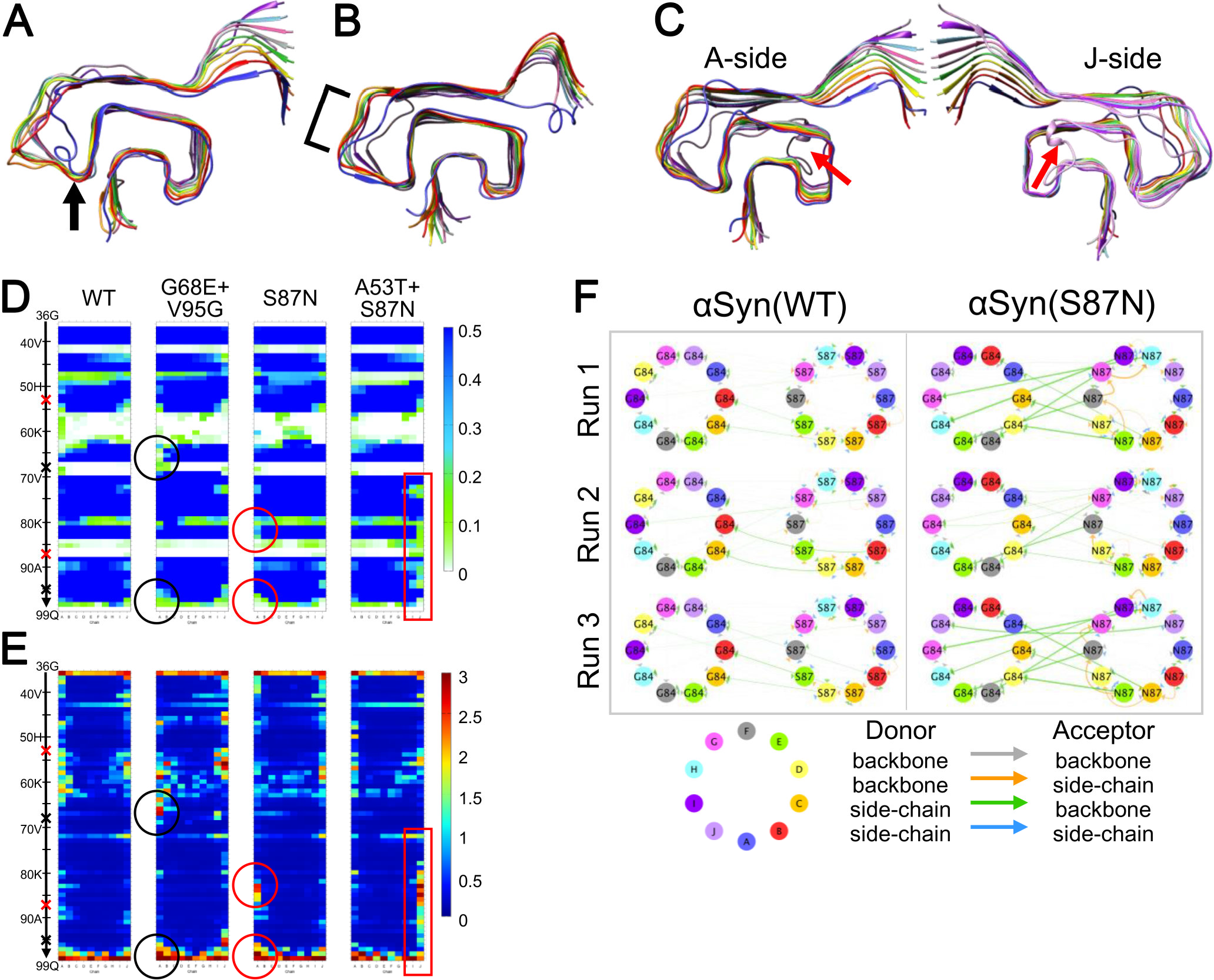
Sheep- or mouse-αSyn-derived mutations which reportedly affect cross-seeding with human αSyn exert definite but limited effects under the simulation condition. **A-C.** Representative images of the homo-(G68E+V95G) (**A**), homo-S87N (**B**) and home-(A53T+S87N) (**C**) amyloids after 400 ns of simulation. All the images from the three independent runs for each are provided in the **Supplementary Figure (Suppl Fig S11B).** As for the homo-(A53T+S87N) amyloid, image viewed from the Chain-J side is also presented, because it reproducibly exhibited severe instability in the β4-β7 region of the Chain J (red arrows), whereas the β7 region of the Chain A was relatively stable despite the A53T substitution. **D-E.** Heatmaps of the β-sheet propensities (**D**) and backbone local B-factors (**E**) of WT, homo-(G68E+V95G), homo-S87N and home-(A53T+S87N) amyloids. The values are average of three runs for the mutants and five runs for WT. All the images from the three independent runs for each are provided in the **Supplementary Figure (Suppl Fig S11C-D).** Comparison of those parameters showed that the structural derangement were limited to the vicinity of each mutation, in one to two stack-end layers **(black circles, red circles, and red box).** The red crosses and the black crosses on the vertical axis represent the positions of mutations of αSyn(A53T+S87N) and αSyn(G68E+V95G), respectively. **F**. Diagrams illustrating hydrogen-bonds between Gly84 and Ser87, comparing those of WT and homo-S87N amyloids. The thickness of each arrow correlates with occupancy of the hydrogen bond. Note that inter-layer bonds are augmented in homo-S87N amyloid; specifically, the side chains of the residues 87 form hydrogen bonds with the backbone of the residues 84 one-layer up, e.g. from the residue 87 of the Chain G to the 84 of the Chain F. These inter-layer bonds could contribute to the stabilization of the loop region in the homo-S87N amyloid. The similar pattern of hydrogen bonds were also seen in the homo-(A53T+S87N) amyloid: All the diagrams of the five runs of WT and the three runs of homo-(A53T+S87N) amyloids are provided in **Supplementary Figure (Suppl Fig S10).**

Luk et al. systematically created various human-mouse chimeric αSyn and investigated their cross-seeding efficiencies by *in vitro* fibril formation and *in vivo* propagation in mice [9] and demonstrated that A53T, S87N and their combination differentially affect the cross-seeding efficiencies. In MD simulation of the mutant αSyn with S87N substitution alone [αSyn(S87N)], the homo-oligomer amyloid of αSyn(S87N) (homo-S87N) seemed rather stable with apparent stabilization of β3, β4, and the intervening loop (**Fig 8A, middle, bracket**). This was interesting because Asn87 cannot directly interact with those regions nor β5 through which β3 could be indirectly affected. We analyzed hydrogen bonds and found that an Asn87 formed an inter-layer hydrogen bond with the carbonyl oxygen of the Gly84 of one-up layer, e.g. the Asn87 of the Chain D and the Gly84 of the Chain C (**Fig 8D**). Those interactions across the loop encompassing 84-87 probably stabilize the local structures around the loop including β6 and proximal β7, and their stabilization could subsequently stabilize the interacting/flanking regions like β2, β3 and distal β5, and so on.

MD simulation of mutant αSyn combining A53T and S87N [αSyn(A53T+S87N)], which has the identical primary structure with the residues 36-99 of mouse αSyn, was completely unexpected. The β5-7 regions of the Chain J were so severely disordered as to dissociate from the stack (**Fig 8A, red arrows**), while the Chain-A side was rather calm (**Fig 8A, right “A-side”**). Correspondingly, β-sheet propensity values of the Chain J were extensively diminished from the β5 to the β7 regions (**Fig 8B, red box**) and the local B-factor values in the region were also high (**Fig 8C, red box**). Possibly, the structural strains posed by A53T substitution, which manifested in β7 of the Chains A and B in the home-A53T, were “redirected” toward the Chain J in the homo-(A53T+S87N) amyloid due to the inter-layer interactions by S87N substitution fixating the loop 84-87 (**Suppl Fig S10**). Since inefficient cross-seeding of mouse αSyn by human αSyn preformed fibrils indicates that the conformation of the human αSyn amyloid is not very compatible with mouse αSyn [9], the drastic destabilization of the Chain J of homo-(A53T+S87N) amyloid might correspond to the incompatibility of mouse αSyn observed in the experiments. Whether the mutually distinct properties of homo-A53T, homo-S87N, and homo-(A53T+S87N) amyloids in the MD simulation milieu have biological significance is very intriguing and to be investigated in the future.

## Discussions

The MD simulation of WT and mutant αSyn amyloids successfully revealed unique intrinsic properties of each which possibly affect amyloid formation reactions. One challenge was the subtleness of the structural alterations caused by mutations under the conditions of MD simulation. Even the stacks of αSyn(G68E+V95G) or αSyn(A53T+S87N), which are reportedly inefficiently cross-seeded by human αSyn amyloids and should be relatively incompatible with the conformation [40][39], showed the unexpectedly small structural aberrations in the limited sampling time frame. Regardless, by repetition of runs to circumvent stochastic factors, certain signs of the incompatibility manifested as substantial local destabilization in the relevant regions. From this view point, the unique destabilization of β7 over two layers, the Chains A-B, by A53T seemed to be consistent as a manifestation of substantial incompatibility with the Greek-key conformation. Another limitation is that the investigated proteins are “artificial proteins” derived from human αSyn by truncation of the unstructured N-terminal and the C-terminal moieties. Since those regions could contribute to amyloid-formation efficiency or strain-specific properties of αSyn amyloids [14][9], the findings on the truncated molecules cannot be directly extrapolated to the actual αSyn amyloids observed *in vitro* or *in vivo.* We think that the present model is still legitimate as a structural model of an in-register parallel β-sheet amyloid in a Greek-key conformation, and that identification of the most incompatible regions of mutant amyloids would be informative about their preferred/compatible amyloid forms. Actual biological relevance of the findings need to be ascertained by *in vitro* experiments with recombinant proteins in the future.

## Asymmetry between the Chain-A and the Chain-J sides of the stack

MD simulation proved to be particularly advantageous in investigations of behaviors of the stack-end molecules, e.g. Chain A and J. They should be important for properties of the amyloid like growth rates of the amyloid fibrils and even the repertoire of non-amyloid proteins they interact, because they are the interfaces which directly interact with the cognate proteins or non-amyloidogenic proteins [41]. Since the behaviors of the stack-end molecules are not recognizable to conventional structural analysis methods, MD simulation can greatly contribute to amyloid researches. A noticeable finding here regarding the stack-end molecules was the apparent asymmetry in structural stabilities between Chain-A and Chain-J sides of the stacks, which reminded that elongation of αSyn fibrils often shows directionality [42]. What mechanism underlies the asymmetry? One possibility is twisting of parallel β-sheets. MD simulation of the αSyn amyloids demonstrated that all the β-sheets incessantly moved in a vibratory manner in narrow spaces, except for β1 and β2 which tended to freely twist and fan (**Supplementary Movie**). Considering the intrinsic twisting tendency [43], it is conceivable that the other β-sheets also share the tendency to twist like β1 and β2. Although they are not allowed to twist freely due to the compact packaging in the Greek-key conformation, even small twists can cause differences in the distances of β-sheets between on the Chain-A side and on the Chain-J side, and consequently differences in structural stabilities because hydrogen bonds or hydrophobic effects are greatly affected by the distances. Another possibility stems from a fact that all the β-strands of a layer of the stack are not necessarily on the same flat plane, as seen in a tau amyloid [8] or an Aβ amyloid [5]. In the case, inter-layer interactions of the β-strands can contribute to the asymmetry because obviously the stack-end molecules lack the interactions in distinct ways. In this regard, the severe destabilization of the Chain J of the homo-(A53T+S87N) is interesting because both the A53T and the S87N mutations newly formed inter-layer hydrogen bonds, which possibly modulate “vertical” positional relations and/or the twisting tendencies of the bonded β-sheets. Whether the directionality of amyloids contribute to cross-seeding/strain barriers is to be investigated.

## Remote effects of the mutations

The contrasting remote influences of A53T and G51D on the stability of the β7 region, in spite that both the mutations are in the β3 region, were most interesting in this study. Given the existence of hydrogen bonds and hydrophobic effects between β3, β5 and β7 as revealed by the MD simulation, alterations in those interactions seem the most likely mechanism of the remote effects. In addition, the intrinsic twisting tendency of β-sheets could modulate the interactions because those properties of β-sheets affect each other [43]. The extra inter-layer hydrogen bonds of homo-A53T amyloid between Thr53 and Gly73 and/or Val74 suggest more intimate β3-β5 interactions (**Fig 3**), which also explain the alterations of β-sheet propensities in the residues of β5. In contrast to the augmented β3-β5 interactions in homo-A53T amyloid, the interactions were obviously disrupted in homo-G51D amyloid due to the severely-disordered β3. The relatively stable β7 regions of homo-G51D amyloid suggest that the disruption of the β3-β5 interactions is advantageous for the structural stability of β7, as if the three β-sheets of the WT amyloid are in balance, i.e. β3-β5 vs β5-β7. A mutation in a Greek-key-type amyloid can affect structural stability of a remote region through successive interactions between β-sheets; this notion could be generalized to any amyloid with a similar architecture.

## Multiple interfaces/amyloid cores and strain diversity

We think that the variable stability of the β7 region of αSyn depending on the intimacy of β3-β5 interactions exemplifies an underlying mechanism of phenotypic diversity of amyloids, or prion strains. If each of β3-β5 and β5-β7 pairs independently functions as an interface/amyloid core of the amyloid, a substrate αSyn which favors intimate β3-β5 interactions would be preferentially incorporated into the amyloids with predominant β3-β5 amyloid core; oppositely, αSyn with inefficient β3-β5 interactions would be preferentially incorporated in the amyloids with predominant β5-β7, because blocking the pathway would facilitate reactions to the other pathway. Those two types of amyloids are therefore equivalent to two different “strains” with respective unique substrate preferences. From this view point, homo-A53T and homo-G51D amyloids are equivalent to the αSyn amyloids with β3-β5 amyloid core and β5-β7 core, respectively (**Fig 9, B and C**). Indeed, amyloids of αSyn(A53T) and αSyn(G51D) are likely to be different “strains” because of the clinicopathological pictures of the associated familial PD [10] and cross-seeding properties *in vitro* [44]. Detailed structures of their amyloid forms are unidentified yet, but the destabilization patterns observed in the MD simulations are informative of their preferred conformations. Considering that amyloid formation of those mutant αSyn are not inhibited by the coexisting αSyn(WT) in the heterozygous familial-PD patients, conformations of the pathogenic amyloids of αSyn(A53T) or αSyn(G51D) could be ones which reconcile the preferences of αSyn(WT) and those mutants.

**Fig 9.**
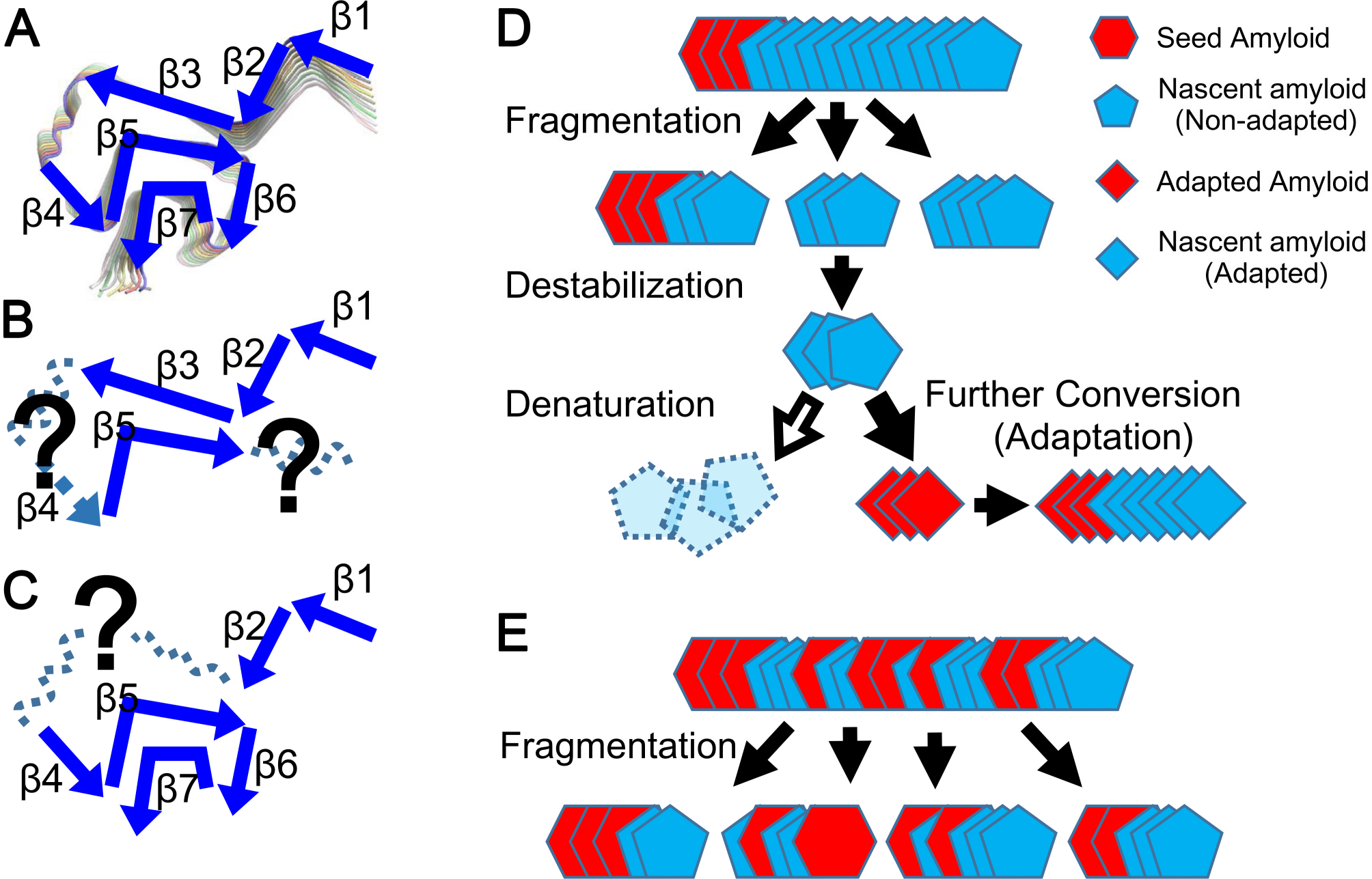
Schematic illustrations: A possible mechanism of strain propensity of αSyn amyloids, and what occurs in a cross-seeding reaction where unstable homo-oligomer and stable hetero-oligomer amyloids are produced. **A.** The WT amyloid has balanced β3, β5 and β7, to maintain the conformation. **B-C.** In an αSyn amyloid with more intimate β3-β5 interactions, like homo-A53T amyloid, the β7 region is destabilized and the conformation would be inclined toward a distinct one compatible with the stable β3-β5 structures **(B)**. To the contrary, in an αSyn amyloid with less β3-β5 interactions, like G51D amyloid, the β7 region is more stabilized and the resultant amyloid would consist mainly of the “NAC” region **(C)**. Since preferences for substrates would be distinct between those two types of amyloids, they can be regarded as two different strains. **D-E.** Schematic illustrations postulating how stable hetero-oligomer and unstable homo-oligomer amyloids can affect the propagation in cross-seeding reactions, or heterologous transmissions of prions. **D.** When seeds are present in an environment where heterologous substrates alone exist, e.g. interspecies transmission of prions, initially the stable hetero-oligomers can grow by incorporating the heterologous substrates, even if the conformation might not be completely the same as the original (top). However, after fragmentation of the grown amyloids, some still remain as stable hetero-oligomers containing the original nuclei, whereas others lose the original nuclei to become unstable homo-oligomers (upper middle). If the destabilized homo-oligomer lose the templating activity (white arrow), the oligomers cannot grow and are eventually degraded. On the other hand, if the destabilized homo-oligomers acquire a novel stable conformation suitable for the substrate along with templating activity, the new seeds would efficiently propagates, emerging as an “adapted strain”. **E.** When seeds are present in an environment where both homologous and heterologous substrates coexist, e.g. a host heterozygous for a pathogenic mutation in an amyloidogenic protein, the amyloid grows as a hetero-oligomer incorporating both types of substrates (top), and the fragmentation produces mainly hetero-oligomers which maintain the templating activity (bottom) to keep propagating as hetero-oligomer amyloids.

The balanced three β-sheets, β3, β5 and β7, are characteristic of WT amyloid: possibly αSyn(WT) might be less efficient to form aggregates than αSyn(A53T) or αSyn(G51D) because it has to reconcile β3-β5 and β5-β7 interactions, unlike the mutants. If the balances between the β-sheets of the WT amyloids could be deranged even without mutations and the amyloid switches the phenotype to ones like homo-A53T or like homo-G51D amyloid, it could explain existence of multiple conformations of αSyn(WT) amyloids, e.g. “ribbon” and “string” which share many β-sheets [45], and also explain why αSyn(WT) is efficiently seeded by either preformed fibrils of αSyn(A53T) or αSyn(G51D) depending on conditions [42][44]. This modality of mechanism for strain diversity can generally occur to parallel β-sheet amyloids with more than two β-sheets aligned side-by-side, and can be operative also in some phenomena about prions (**Supplementary Discussion**).

## Effects of hetero-oligomerization

The remote effects of the A53T substitution on the stability of β7 was unexpected, and the suppression of the unique influences of A53T by hetero-oligomerization with the αSyn(WT) was another surprise. As discussed earlier, the unique pattern of destabilization of β7 of homo-A53T amyloid as indicated by the low β-sheet propensity along with the large local B-factor values suggested incompatibility of αSyn(A53T) with the Greek-key conformation. αSyn(A53T) may actually prefer conformations with more intimate β3-β5 interactions, e.g. the speculative model postulated by Rodriguez et al. where the “pre-NAC” region containing Thr53 and the “NACore” around Gly73 tightly bind [37]. It is also reported that *in vitro*-formed αSyn(WT) fibrils and αSyn(A53T) fibrils are morphologically distinct [42]. The signs of the incompatibility of αSyn(A53T) with the Greek-key conformation was absent in any hetero-A53T amyloids that we tested, i.e. they were suppressed just by hetero-oligomerization with αSyn(WT). This notion helps address the question why A53T cause familial PD despite the incompatibility of αSyn(A53T) with the Greek-key conformation, without interfering with amyloid formation (**Fig 9C and D**). On the other hand, the destabilization of β7 region observed in hetero-H50Q(ABCDE) demonstrated that hetero-oligomerization with αSyn(WT) can destabilize the amyloid stack depending on the mutation types. The MD simulation of the hetero-oligomers also implied a possibility that they can combine properties of WT and the mutant, e.g. higher β-sheet propensity values in the β3 of hetero-A53T amyloids than WT. Those notions need to be considered in interpretation of cross-seeding and inter-species transmission of prions (**Supplementary Discussion**). Whether stability of hetero-oligomers corresponds to cross-reaction efficiency also needs further investigation in the future.

In summary, the present series of MD simulations of αSyn amyloids revealed the following: (1) structural asymmetry between the Chain-A and the Chain-J sides, (2) a mutation can affect stabilities of remote β-sheets through complex interactions between β-sheets, (3) existence of at least two potential interfaces/amyloid cores that can be switched by mutations, (4) the switchable interfaces possibly underlie the multiple αSyn-amyloid strains, and (5) unique properties of the homo-oligomer amyloid of mutant αSyn can be obscured by hetero-oligomerization with the wild-type. It was surprising that a stack of only ten peptides with moderately intricate conformation exhibit those properties reminiscent of complex phenomena about prions as discussed in detail in the **Supplementary Discussion**. Those findings support our view that prion-like properties are inherent in the in-register parallel β-sheet structures. Thus, *in silico* studies are potentially powerful investigation tools of in-register parallel β-sheet amyloids. The present studies are valuable example of MD simulation of this type of amyloids with a relatively large size and complex architecture. Hopefully the knowledge from the studies are applicable to other amyloids including prion, Aβ peptide, and tau, and contribute to advance our understanding of their pathogenicity and pathophysiology.

## Acknowledgement

The numerical calculations were carried out on the TSUBAME2.5/3.0 supercomputer in the Tokyo Institute of Technology and Reedbush-U supercomputer in the Information Technology Center, The University of Tokyo. This work is supported by “TSUBAME Encouragement Program for Young/Female Users” of Global Scientific Information and Computing Center at Tokyo Institute of Technology, “Initiative on Promotion of Supercomputing for Young or Women Researchers” of Information Technology Center, The University of Tokyo, and Takeda Science Foundation.

## References

1. Prusiner SB. Prions. Proc Natl Acad Sci U S A. 1998;95: 13363–13383.

2. Telling GC, Parchi P, DeArmond SJ, Cortelli P, Montagna P, Gabizon R, et al. Evidence for the conformation of the pathologic isoform of the prion protein enciphering and propagating prion diversity. Science. 1996;274: 2079–82. Available: http://www.ncbi.nlm.nih.gov/pubmed/8953038

3. Groveman BR, Dolan MA, Taubner LM, Kraus A, Wickner RB, Caughey B. Parallel in-register intermolecular β-sheet architectures for prion-seeded prion protein (PrP) amyloids. J Biol Chem. 2014;289: 24129–24142. doi:10.1074/jbc.M114.578344

4. Vázquez-fernández E, Vos MR, Afanasyev P, Cebey L, Sevillano AM, Vidal E, et al. The Structural Architecture of an Infectious Mammalian Prion Using Electron Cryomicroscopy. PLoS Pathog. 2016;12: e1005835. doi:10.1371/journal.ppat.1005835

5. Schmidt M, Rohou A, Lasker K, Yadav JK, Schiene-fischer C, Fändrich M, et al. Peptide dimer structure in an Aβ(1-42) fibril visualized with cryo-EM. Proc Natl Acad Sci U S A. 2015;112: 11858–11863. doi:10.1073/pnas.1503455112

6. Gremer L, Schölzel D, Schenk C, Reinartz E, Labahn J, Ravelli RBG, et al. Fibril structures of amyloid-β(1-42) by cryoelectron microscopy. Science. 2017;358: 116–119.

7. Tuttle MD, Comellas G, Nieuwkoop AJ, Covell DJ, Berthold DA, Kloepper KD, et al. Solid-state NMR structure of a pathogenic fibril of full-length human α-synuclein. Nat Struct Mol Biol. 2016;23: 409–415. doi:10.1038/nsmb.3194

8. Fitzpatrick AWP, Falcon B, He S, Murzin AG, Murshudov G, Garringer HJ, et al. Cryo-EM structures of tau filaments from Alzheimer’s disease. Nature. Nature Publishing Group; 2017;547: 185–190. doi:10.1038/nature23002

9. Luk KC, Covell DJ, Kehm VM, Zhang B, Song IY, Byrne MD, et al. Molecular and Biological Compatibility with Host Alpha-Synuclein Influences Fibril Pathogenicity. Cell Rep. 2016;16: 3373–3387. doi:10.1016/j.celrep.2016.08.053

10. Petrucci S, Ginevrino M, Valente EM. Parkinsonism and Related Disorders Phenotypic spectrum of alpha-synuclein mutations◻: New insights from patients and cellular models. Park Relat Disord. Elsevier Ltd; 2016;22: S16–S20. doi:10.1016/j.parkreldis.2015.08.015

11. Kim WS, Kågedal K, Halliday GM. Alpha-synuclein biology in Lewy body diseases. Alzheimers Res Ther. 2014;6: 73.

12. Bisaglia M, Trolio A, Bellanda M, Bergantino E, Bubacco L, Mammi S. Structure and topology of the non-amyloid-β component fragment of human α-synuclein bound to micelles◻: Implications for the aggregation process. Protein Sci. 2006;15: 1408–1416. doi:10.1110/ps.052048706.Alzheimer

13. Lashuel HA, Overk CR, Oueslati A, Masliah E. The many faces of α◻synuclein◻: from structure and toxicity to therapeutic target. Nat Rev Neurosci. Nature Publishing Group; 2013;14: 38–48. doi:10.1038/nrn3406

14. Masuda-suzukake M, Nonaka T, Hosokawa M, Oikawa T, Arai T, Akiyama H, et al. Prion-like spreading of pathological α-synuclein in brain. Brain. 2017;136: 1128–1138. doi:10.1093/brain/awt037

15. Shimozawa A, Ono M, Takahara D, Tarutani A, Imura S, Masuda-suzukake M, et al. Propagation of pathological α-synuclein in marmoset brain. Acta Neuropathologica Communications; 2017; 1–14. doi:10.1186/s40478-017-0413-0

16. Xu L, Ma B, Nussinov R, Thompson D. Familial Mutations May Switch Conformational Preferences in α◻ Synuclein Fibrils. ACS Chem Neurosci. 2017;8: 837–849. doi:10.1021/acschemneuro.6b00406

17. Case DA, Betz RM, Cerutti DS, Cheatham, III TE, Darden TA, Duke RE, et al. AmberTools16. University of California, San Francisco. 2016. doi:10.1021/ct200909j

18. Krivov GG, Shapovalov M V, Dunbrack RL. Improved prediction of protein side-chain conformations with SCWRL4. Proteins Struct Funct Bioinforma. Wiley Subscription Services, Inc., A Wiley Company; 2009;77: 778–795. doi:10.1002/prot.22488

19. Abraham MJ, Murtola T, Schulz R, Páll S, Smith JC, Hess B, et al. GROMACS: High performance molecular simulations through multi-level parallelism from laptops to supercomputers. SoftwareX. 2015;1-2: 19–25. doi:https://doi.org/10.1016/j.softx.2015.06.001

20. Lindorff-Larsen K, Piana S, Palmo K, Maragakis P, Klepeis JL, Dror RO, et al. Improved side-chain torsion potentials for the Amber ff99SB protein force field. Proteins Struct Funct Bioinforma. Wiley Subscription Services, Inc., A Wiley Company; 2010;78: 1950–1958. doi:10.1002/prot.22711

21. Jorgensen WL, Chandrasekhar J, Madura JD, Impey RW, Klein ML. Comparison of simple potential functions for simulating liquid water. J Chem Phys. American Institute of Physics; 1983;79: 926–935. doi:10.1063/1.445869

22. Bussi G, Donadio D, Parrinello M. Canonical sampling through velocity rescaling. J Chem Phys. American Institute of Physics; 2007;126: 14101. doi:10.1063/1.2408420

23. Berendsen HJC, Postma JPM, van Gunsteren WF, DiNola A, Haak JR. Molecular dynamics with coupling to an external bath. J Chem Phys. American Institute of Physics; 1984;81: 3684–3690. doi:10.1063/1.448118

24. Hess B. P-LINCS:◻ A Parallel Linear Constraint Solver for Molecular Simulation. J Chem Theory Comput. American Chemical Society; 2008;4: 116–122. doi:10.1021/ct700200b

25. Essmann U, Perera L, Berkowitz ML, Darden T, Lee H, Pedersen LG. A smooth particle mesh Ewald method. J Chem Phys. American Institute of Physics; 1995;103: 8577–8593. doi:10.1063/1.470117

26. Páll S, Hess B. A flexible algorithm for calculating pair interactions on SIMD architectures. Comput Phys Commun. 2013;184: 2641–2650. doi:https://doi.org/10.1016/j.cpc.2013.06.003

27. Fuchs JE, Waldner BJ, Huber RG, von Grafenstein S, Kramer C, Liedl KR. Independent Metrics for Protein Backbone and Side-Chain Flexibility: Time Scales and Effects of Ligand Binding. J Chem Theory Comput. American Chemical Society; 2015;11: 851–860. doi:10.1021/ct500633u

28. Roe DR, Cheatham TE. PTRAJ and CPPTRAJ: Software for Processing and Analysis of Molecular Dynamics Trajectory Data. J Chem Theory Comput. American Chemical Society; 2013;9: 3084–3095. doi:10.1021/ct400341p

29. Humphrey W, Dalke A, Schulten K. VMD: Visual molecular dynamics. J Mol Graph. 1996;14: 33–38. doi:https://doi.org/10.1016/0263-7855(96)00018-5

30. Touw WG, Baakman C, Black J, te Beek TAH, Krieger E, Joosten RP, et al. A series of PDB-related databanks for everyday needs. Nucleic Acids Res. Oxford University Press; 2015;43: D364–D368. doi:10.1093/nar/gku1028

31. Kabsch W, Sander C. Dictionary of protein secondary structure: Pattern recognition of hydrogen-bonded and geometrical features. Biopolymers. Wiley Subscription Services, Inc., A Wiley Company; 1983;22: 2577–2637. doi:10.1002/bip.360221211

32. Tiberti M, Invernizzi G, Lambrughi M, Inbar Y, Schreiber G, Papaleo E. PyInteraph: A Framework for the Analysis of Interaction Networks in Structural Ensembles of Proteins. J Chem Inf Model. American Chemical Society; 2014;54: 1537–1551. doi:10.1021/ci400639r

33. Salamanca Viloria J, Allega MF, Lambrughi M, Papaleo E. An optimal distance cutoff for contact-based Protein Structure Networks using side-chain centers of mass. Sci Rep. England; 2017;7: 2838. doi:10.1038/s41598-017-01498-6

34. Shannon P, Markiel A, Ozier O, Baliga NS, Wang JT, Ramage D, et al. Cytoscape: A Software Environment for Integrated Models of Biomolecular Interaction Networks. Genome Res. 2003;13: 2498–2504. doi:10.1101/gr.1239303

35. Pettersen EF, Goddard TD, Huang CC, Couch GS, Greenblatt DM, Meng EC, et al. UCSF Chimera—A visualization system for exploratory research and analysis. J Comput Chem. John Wiley & Sons, Inc.; 2004;25: 1605–1612. doi:10.1002/jcc.20084

36. Fuchs JE, Waldner BJ, Huber RG, Grafenstein S Von, Kramer C, Liedl KR. Independent Metrics for Protein Backbone and Side-Chain Flexibility: Time Scales and E ff ects of Ligand Binding. J Chem Theory Comput. 2015;11: 851–60. doi:10.1021/ct500633u

37. Rodriguez JA, Ivanova MI, Sawaya MR, Cascio D, Reyes FE, Shi D, et al. Structure of the toxic core of α-synuclein from invisible crystals. Nature. 2015;525: 486–90. doi:10.1038/nature15368

38. Blauwendraat C, Kia DA, Pihlstrøm L, Gan-Or Z, Lesage S, Gibbs JR, et al. Insufficient evidence for pathogenicity of SNCA His50Gln (H50Q) in Parkinson’s disease. Neurobiol Aging. Elsevier Inc; 2017;64: 159.e5–159.e8. doi:10.1016/J.NEUROBIOLAGING.2017.12.012

39. Bickle L, Hopwood JJ, Karageorgos L. Analysis of sheep α-synuclein provides a molecular strategy for the reduction of fibrillation. BBA - Proteins Proteomics. Elsevier B.V.; 2017;1865: 261–273. doi:10.1016/j.bbapap.2016.12.008

40. Moriarty GM, Olson MP, Atieh TB, Janowska MK, Khare SD, Baum J. A pH-dependent Switch Promotes α-synuclein Fibril Formation via Glutamate Residues. J Biochem. 2017;292: 16368–16379. doi:10.1074/jbc.M117.780528

41. Richardson JS, Richardson DC. Natural beta-sheet proteins use negative design to avoid edge-to-edge aggregation. Proc Natl Acad Sci U S A. 2002;99: 2754–9. doi:10.1073/pnas.052706099

42. Sidhu A, Segers-Nolten I, Subramaniam V. Conformational Compatibility Is Essential for Heterologous Aggregation of α-Synuclein. ACS Chem Neurosci. 2016;7: 719–27. doi:10.1021/acschemneuro.5b00322

43. Periole X, Huber T, Bonito-Oliva A, Aberg KC, van der Wel PCA, Sakmar TP, et al. Energetics Underlying Twist Polymorphisms in Amyloid Fibrils. J Phys Chem B. 2018;122: 1081–1091. doi:10.1021/acs.jpcb.7b10233

44. Sierecki E, Giles N, Bowden Q, Polinkovsky ME, Steinbeck J, Arrioti N, et al. Nanomolar oligomerization and selective co-aggregation of α-synuclein pathogenic mutants revealed by single-molecule fluorescence. Sci Rep. Nature Publishing Group; 2016;6: 37630. doi:10.1038/sreβ37630

45. Meier BH, Gath J, Bousset L, Habenstein B, Melki R, Bo A. Unlike Twins◻: An NMR Comparison of Two α-synuclein Polymorphs Featuring Different Toxicity. PLoS One. 2014;9: e90659. doi:10.1371/journal.pone.0090659

